# Promyelocytic Leukemia Protein regulates Angiogenesis and Epithelial-Mesenchymal Transition to limit metastasis in MDA-MB-231 breast cancer cells

**DOI:** 10.1101/2022.08.28.505601

**Authors:** Amalia P Vogiatzoglou, Syrago Spanou, Nikoleta Sachini, Elias Drakos, Christoforos Nikolaou, Takis Makatounakis, Androniki Kretsovali, Joseph Papamatheakis

## Abstract

Promyelocytic Leukemia Protein (PML) is the core protein of nuclear bodies (NBs) that regulate a large number of cellular processes, including, context dependent, tumor –suppressor and pro-oncogenic effects. PML knockdown (KD) in breast cancer lines MDA-MB-231, but not MCF7, cells showed higher cell proliferation, increased migration properties and prolonged stem cell –like survival in line with gene enrichment results from RNA sequencing analysis. MDA-MB-231 PML KD cells showed an increase of hypoxic and mesenchymal characteristics manifested by higher HIF1a and the EMT-TWIST2 protein levels respectively. Mechanistically, PML loss caused an increase of HIF1a and TWIST2 RNA levels. Interestingly, TWIST2 binds to PML. Moreover, PML directly opposed the action of HIF1a and TWIST2 on VEGFa and CD24 reporters, respectively. Tumor xenografts of MDA-MB-231 PML KD cells showed a higher micro vessel content, grew faster and had higher metastatic ability with a preference for lung. Thus, PML opposes the aggressive cancer phenotype by multiple mechanisms that antagonize the HIF-hypoxia-angiogenic and EMT axis.

## 1. INTRODUCTION

Breast cancer is characterized by genetic and epigenetic complexity and phenotypic heterogeneity[1]. Differences between breast cancer-subtypes or patients referred as intertumoral heterogeneity as well as cellular diversity within the same tumors[2] is dynamically driven by genetic and epigenetic evolution and has an impact in patient’s prognosis and therapy. Major current efforts are directed towards the elucidation of factors that modify the action of critical oncogene or tumor suppressor pathways that regulate invasiveness, metastatic potential and drug response with predictive or therapeutic importance.

The Promyelocytic Leukemia (PML) gene was first described in the early 1990s at the point of chromosomal translocation t (15, 17), where it was found to encode an oncogenic chimeric protein emerging from the fusion of PML and the retinoic acid receptor alpha (RAR-α) in patients with acute promyelocytic leukemia (APL). The PML-RAR chimeric protein prevents the differentiation of bone marrow progenitor cells, sustaining their self-renewal capacity and preventing their apoptotic process, leading to leukemogenesis[3,4]. Promyelocytic leukemia protein is the key component of PML-NBs, which are nuclear membrane less compartments self-assembled through the RBCC motif interactions. By interacting and/or participating in the post-translational modifications of various nuclear proteins, PML-NBs regulate a variety of cellular processes such as apoptosis, proteolysis, cellular aging, cell self-renewal capacity, DNA damage response, telomere stability, gene expression and chromatin/epigenetic states[5,6]. For example PML is highly expressed during murine mammary gland development, it’s barely detectable during pregnancy and lactation and it’s necessary for lineage commitment of bi-potent luminal progenitor cells[7]. Moreover PML is essential for sustaining naïve pluripotent state in mouse embryonic stem cells[8].The role of PML in neoplasias seems to be complex. PML protein is lost in human cancers of various histologic origins, including breast cancer[9]. Various reports support a tumor suppressive role for PML[10]. Using an inducible expression of PML IV in the triple negative (TNBC) Claudin low (CLD) subtype MDA-MB-231 cell line, we found strong inhibition of cell proliferation and tumor sphere formation in a reversible manner[11]. Other studies point to a pro-survival role of PML in non-tumorigenic or tumorigenic breast cell lines and find that high PML levels correlate with poor prognosis in a breast cancer cohort[12]. Similarly, high PML expression in chronic myeloid leukemia maintains hematopoietic stem cells and PML ablation leads to an increase of their initial cycling activity that eventually leads to their eradication[13]. These studies suggest that PML may have divergent cells effects not only in different cell contexts but also in different temporal windows i.e. short vs long-term effects.

Here we show that constitutive PML loss enhances cell migration *in vitro* & metastatic aggressiveness *in vivo* of TNBC cells. Cells derived from *in vivo* primary tumors or lung metastasis stably maintain an *in vivo* aggressive phenotype and transcriptionally resemble the earlier described lung metastasis signature (LMS)[14] that may result from either epigenetic or clonal evolution during *in vivo* growth in the absence of PML. Interestingly, we found that PML interacts and likely functionally affects the activity of various bHLH or Zinc finger type proteins including the master hypoxia regulator HIF1a and EMT mediators, such as TWIST2. Thus, PML may impede the aggressive EMT-metastatic behavior of TNBC cells, in line with transcriptional profiling that shows enrichment in adhesion, cell cycle and signaling categories deregulated by PML ablation.

## 2. MATERIALS AND METHODS

### 2.1. Cell culture and generation of stable cell lines

The 293T cell line was obtained from the American Type Culture Collection (ATCC, Manassas,VA,USA), MCF7 and MDA-MB-231 from the Deutsche Sammlung von Mikroorganismen und Zellkulturen GmbH (DSMZ, Braunschweig – Germany). They were authenticated in the past 2 years by PCR-single locus technology (Eurofins Genomics-Europe Applied Genomics GmbH, Ebersberg, Germany) using the DSMZ (http://www.dsmz.de/de/service/services) and the Cellosaurus database (https://web.expasy.org/cellosaurus). Cells were mycoplasma free as determined by PCR (mycoplasma detection kit “Venor GeM classic “MINERVA BIOLABS, Berlin, Germany).

For the generation of stable, knock down cell lines, cells were infected by puromycin or G418 resistance -lentiviral vectors carrying the relevant shRNA sequences, followed by drug selection. The shPML short-hairpin RNA sequences that we used were: Sh0:5’-AGATGCAGCTGTATCCAAG-3’[15], Sh1: 5’-GCTGTATCCAAGAAAGCCA-3’, sh2: 5-CCAACAACATCTTCTGCTCC-3’ were cloned into the lentiviral vector pLKO.1 (Addgene #8453)[16]. Knockdown (KD) was evaluated by western blot and/or mRNA analysis using suitable primers.

### 2.2. Plasmids, DNA, and siRNA transfections

Plasmids with PMLIV, PMLIII and PMLI isoforms have been previously described [17]. For immunoprecipitation and localization experiments the PMLIV, PMLIII and PMLI isoforms were fused to the mRED vector (Clontech, Mountain View, CA, USA). TWIST1, TWIST2, SNAIL1 (Addgene #16225[18]) and SLUG (Addgene #25696) were fused to GFP-C (Clontech). mCherry LEGO-C2 was provided by Dr. K.Weber. The 5xHREVEGFLuc reporter was provided by Dr. S. Simos. The CD24-Luc was constructed by cloning the −1700 to +79bp (PCR amplified using CD24 forward 5’-GAGCTAAAGTGACTGACCTTGAAGGCACAA-3’ and reverse 5’-CGTCTAGCAGGATGCTGGGTGCTTG-3’ primers) of the CD24 promoter upstream of the pGL3luc reporter. Transient transfections were performed using the calcium phosphate method or Lipofectamine 2000 (Thermo Fisher Scientific, Waltham, MA, USA) according to the manufacturer’s instructions. The lentivirus production and infection protocol have been previously described in detail[19].

### 2.3. Tumor Sphere Forming assay

For tumorsphere forming assays cells were cultured in DMEM-F12 1:1 (Gibco, Waltham, Massachusetts, USA) containing B27 (1:50), bFGF (20ng/ml), EGF (20ng/ml) and 0.2% methylcellulose at 1000 cells/ml using in ultra-low attachment plates. After 8 days, spheres of ≥ 70μm size were enumerated and if required were dispersed by accutase and cells were counted for next passage.

### 2.4. RNA extraction and qRT-PCR

Total RNA extraction was performed using Nucleozol (Macherey-Nagel, Germany). Next, 2μg RNA was used to generate library cDNAs using the enzyme M-MuLV Reverse Transcriptase (New England, Biolabs, USA) together with an RNase inhibitor (New England, Biolabs, USA) according to the manufacturer’s protocol. The relative abundance of each gene transcript was measured by qRT-PCR using the dye SYBR Green I (Invitrogen, Waltham, Massachusetts, USA). Relative mRNA expression was calculated after normalization against β-actin or GAPDH levels. Set of primers used for qRT-PCR are listed in Suppl. Materials file.

### 2.5. RNA sequencing

Total RNA was isolated from control cells and PML KD using TRIzol (Invitrogen). For the MDA-MB-231 samples, the mRNA 3’-UTR sequence method was used to create libraries with the QuantSeq 3’mRNA-Seq Library Prep Kit for Ion Torrent-Cat #012. And the sequencing was done with Ion S5, with Ion 540 Reagents. NEBNext Ultra II Directional RNA Library Prep Kit for Illumina-E7760 and NEBNext Poly (A) mRNA Magnetic Isolation Module-E7490 were used for libraries for complete RNA sequencing of MCF7 cells. Sequencing was performed on the NextSeq 500 Illumina with FlowCell High 1×75. HISAT2 version 2.1 (genome mapper)

### 2.6. Analysis of differentially expressed genes

Differential gene expression analysis was performed with MetaseqR. The functional analysis was performed using the web tool gProfileR[20] and Metascape[21]

### 2.7. Western-Blot Analysis

Whole cell lysates were prepared using RIPA cell lysis buffer (25 mm Tris pH 7.6, 150 mm NaCl, 1% NP-40, 1% deoxycholate, 0.1% SDS, 1 mm PMSF) containing protease inhibitor cocktail (Complete; Sigma), and protein concentration was determined by Bradford assay. Equal amounts of cell lysates were subjected to SDS/PAGE, followed by immunoblotting. The primary antibodies used for WB are listed in Suppl. Materials file.

### 2.8. Protein immunoprecipitation (Immunoprecipitation-IP)

Immunoprecipitation of protein complexes was performed, using HEK293T cell extracts following overexpression of specific proteins for possible interaction and prepared by RIPA cell lysis buffer as described above. 200μg of protein extracts were incubated with primary antibody overnight at 4 °C. The following day, 20 uL of protein G beads was added to each sample after washing with IP buffer (25 mm Tris/HCl pH 7.6, 150 mm NaCl), and reactions were incubated at 4 °C for three additional hours. Non-specific proteins were washed away three times with NETN buffer (10mm Tris/HCl pH 8.0, 250mm NaCl, 5mm EDTA, 0.5% NP-40, 1mm PMSF). SDS sample buffer was added, and the samples were boiled prior to SDS/PAGE analysis. Input lanes represent 10% of the lysate used for the IP.

### 2.9. FACs analysis

Flow cytometry was used to evaluate CD24 and CD44 surface expression (FACs calibour by Becton Dickinson (BD) (Franklin Lakes, New Jersey, USA). The cells were detached with trypsin-EDTA and following centrifugation were suspended in PBS-2% FCS-0.1% NaN3 with specific antibodies of isotype controls.

### 2.10. Transwell migration assay

For migration assays, cells were seeded into Millicell, Cell culture 8μm inserts (Merck KGaA, Germany) in FBS-free medium overlaying the lower compartment filled with FBs-containing media. The next day, the upper side cells were scraped away and the insert was fixed with 70% ethanol, stained with crystal violet (Merck) and counted under a microscope. The percentage of cells that has passed through the membrane in relation to the total number of cells seeded on the transwell is indicative of the migrating capacity of the cells.

### 2.11. Immunostaining and microscopy

Cells for immunostaining were cultured on 8-well chamber, removable (80841, Ibidi GmbH, Germany) and fixed in buffered 4% PFA / 1XPBS for 5 minutes at room temperature, permeated with 0.5% Triton X-100/1XPBS for 5 minutes and rinsed repeatedly with 1XPBS. The samples were then incubated with 1% BSA/1XPBS for 1 hour before incubation with primary antibodies overnight or 1 hour. After washing with PBS, secondary antibody was added to the samples for one hour. The secondary antibody was washed again three times with PBS and the cell nuclei were then counter stained with DAPI (Merck). Epifluorescence microscopy was done in an inverted Olympus IX70 and Confocal microscopy was done in a Zeiss Axioscope 2 Plus microscope equipped with a Bio-Rad Radiance 2100 laser scanning system and Lasersharp 2000 imaging software and a Leica SP8 inverted focus microscope and analyzed with the Leica Application Suite (Las, Leica, Germany) software. Autopsied animals were examined under a Leica M205FA fluorescent stereomicroscope, carrying a Leica DFC310FX camera and analysed with Leica Application Suite (Las) software.

### 2.12. Immunohistochemistry in paraffin sections

Tissues isolated from experimental animals were fixed with 10% formalin (Merck) and kept overnight at 4° C. They were then washed 3 times with sterile PBS and placed in 70% sterile ethanol (analytical grade). They were then placed in a sequential dehydration and paraffinization machine and finally the tissues were placed in paraffin to be cut into a microtome and stained with hematoxylin-eosin as described in a previous study[22]. If required, sections were stained with antibody CD31 (AbCAM, Cambridge,UK) and stained area was measured with ImageJ as percentage of Microvessel density.

### 2.13. *In vivo* experiments on experimental animals

Tumor Xenografting was done in NOD-scid IL2R gamma null (NSG) mice. In the case of MCF7 cells, β-estradiol (8μg / ml) was added to the water and was being replaced every 5 days until termination of the experiment. Tumor volume was estimated using the formula V =½ (Length × Width^2^)[23]. To better monitor the engrafted tumour cells, control and KD PML cells were previously modified to express the red fluorescent mCherry protein. Equal cell numbers from either group were injected subcutaneously into NSG mice and measurements from the growing primary tumors were taken every four days. When tumors reached 1200mm^3^ in size, mice were sacrificed.

### 2.14. Approval of animal experiments

All procedures were conducted according to Greek national legislations and institutional policies following approval by the FORTH ethical committee. NSG mice were purchased from Jackson Laboratory (Bar Harbor, Maine, USA) and were maintained and bred at the Institute of Molecular Biology and Biotechnology (IMBB) animal facility following the institutional guidelines based on the Greek ethical committee of animal experimentation. Current procedures were approved by the General Directorate of Veterinary Services, Region of Crete (license number: 93304 and 106336.)

### 2.15. Statistical analysis

The statistics was performed with Microsoft EXCEL software, XLSTAT and Graphpad Prism. Values were presented as the mean ±SD from two or more experiments. For comparisons the two tailed t-test, Mann-Whitney or χ2 tests were performed as descrided in results.

## 3. RESULTS

### 3.1. Silencing of PML affects tumor cell morphology and physiology

To dissect the role of PML in breast cancer we initially employed a Doxycycline inducible shRNA approach that resulted in 50-55% reduction of the various PML isoforms. Thus we switched to a constitutive knock down approach by testing three different shRNAs for PML silencing in MDA-MB-231 and MCF7 cells. Knock down (KD) efficiency was verified by qRTPCR (not shown), immunostaining and western blotting using an antibody against a region common to all PML isoforms (Fig.1A) and cell pools with more than 80% PML protein reduction were chosen for further experimentation.

**Figure 1.**
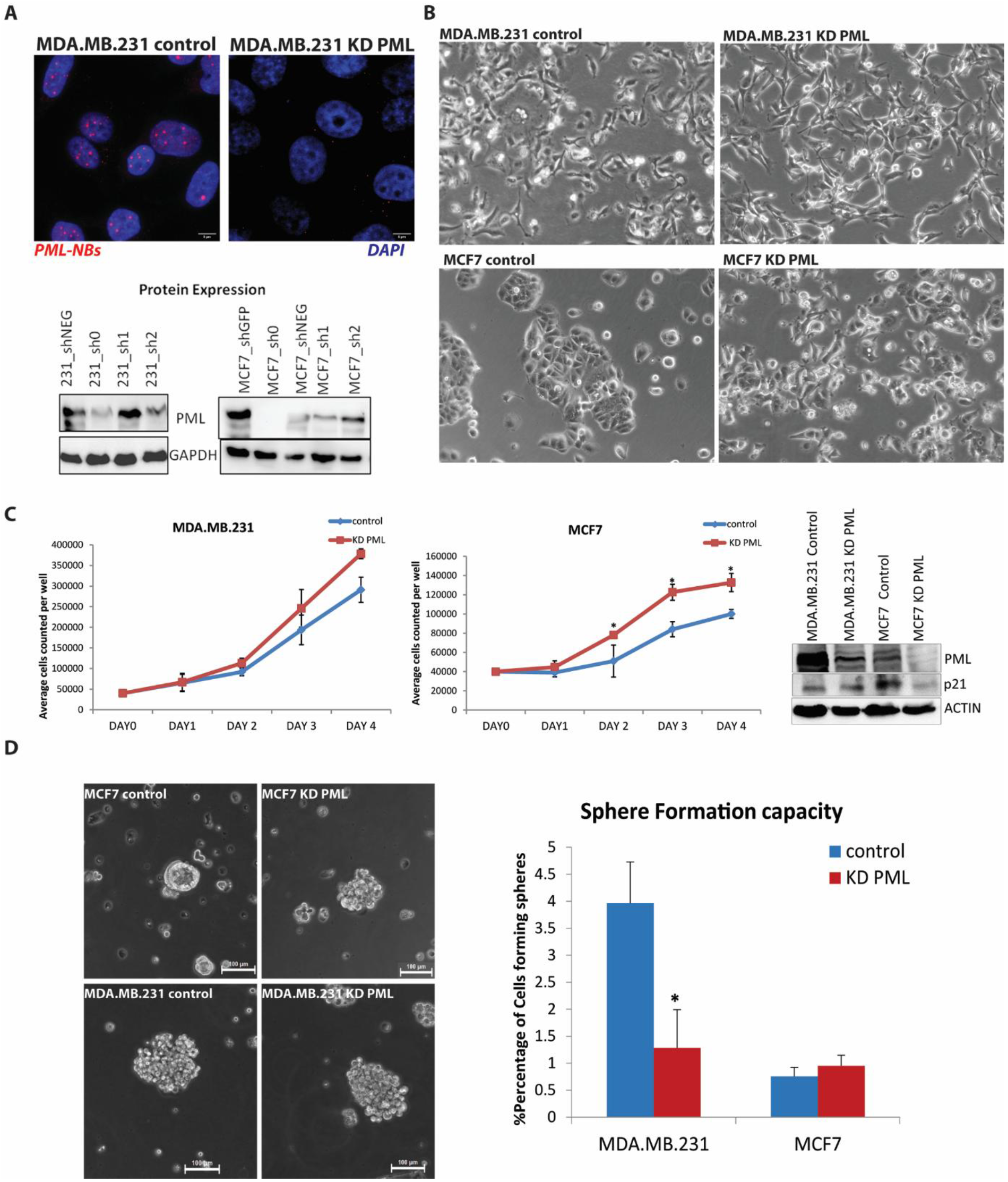
PML silencing alters breast cancer cells phenotype. (A) Upper: Confocal microscopy of PML Immunostaining (red nuclear speckles) in control and PML–KD MDA-MB-231 cells counterstained by DAPI (blue). Lower: Immunoblotting for PML protein expression of puromycin selected pools of MCF7 and MDA-MB-231 using a sh-scrabbled control and 3 different sh-PML (sh0, sh1, sh2) targeting sequences. All further experiments were performed using the most efficient sh0 sequence. (B) Phase contrast microscopy of MDA-MB-231 KD PML & MCF7 KD PML compared to sh-scrabbled controls. (C) Cell growth of control and PML KD, MDA-MB-231 (left) and MCF7 (middle) cells. Results show mean ±SD from one out of three independent triplicate experiments with similar results. t-test, *p-value≤0.05. Right: Immunoblot of p21 protein levels (CDKN1A) in control and PML KD cell lines. (D) Morphology of 3D tumorspheres of control or PML KD, MCF7 and MDA-MB-231 (left) and tumor sphere forming efficiency (right). Shown is mean ±SD of % sphere forming cells from one of at least two quadruplicate experiments with similar results. t-test * p-value ≤0.05

Microscopic examination of control and KD cells showed moderate changes of cell morphology in sparse cultures, towards a more spindle–like shape of MDA-MB-231 (Fig.1B). Similarly, MCF7 acquired a less tight epithelial cell-to-cell adhesion pattern. PML silencing slightly increased the proliferation of MDA-MB-231 and to a larger extend MCF7 in agreement with reduced expression of the cyclin kinase inhibitor p21 protein (Fig.1C). We next tested the impact of PML ablation on cells grown in 3D as tumour spheres. Parental MCF7 and claudin-low MDA-MB-231 formed compact -round and grape-like spheres respectively[24]. The MCF7-PML KD spheres assumed a more loose morphology that tends to resemble those of MDA-MB-231 cells (Fig.1D). Most importantly the sphere forming efficiency of MDA-MB-231 –PML KD cells, but not that of MCF7 cells was reduced to about 30% of the control cells (Fig.1D).

We have shown previously that PML is essential for maintaining the epithelial characteristics of naïve mouse embryonic stem cells[8]. The MDA-MB-231 cell line has mesenchymal characteristics, expressing high levels of VIMENTIN compared to MCF7 (Fig.2A/middle). Conversely, MCF7 express high levels of the epithelial marker CDH1 as compared to MDA-MD-231 cells. Following PML KD we observed a decrease of CDH1 RNA in both MDA–MB-231 and MCF7 cells (Fig 2A/left). At the protein level, the VIMENTIN expression was slightly increased in the former whereas the CDH1 was decreased in the latter cells (Fig.2A/Right). MDA-MB-231 cells express high levels of the mesenchymal marker CD44 and intermediate levels of the epithelial marker CD24, a phenotype early on linked to high tumour initiating (or cancer stem cell) ability of patients-derived breast cancer xenografts[25]. Flow cytometry analysis for those surface markers showed that PML loss minimally affected CD24 or CD44 expression in MCF7 cells. However, PMLKD in MDA-MB-231, led to a reduction of the CD24^+^ population (Fig.2B) and lower CD24 RNA expression (Fig.6D). To further assess the epithelial/mesenchymal cell properties we used an *in vitro* transwell assay, and found that the absence of PML led to increased migration capacity of MDA-MB-231 and to a lesser extend MCF7 cells (Fig.2C). Moreover the amount of phosphorylated STAT3 protein, known to correlate with TNBC aggressive behaviour[26,27], was increased (Fig.2D).

**Figure 2.**
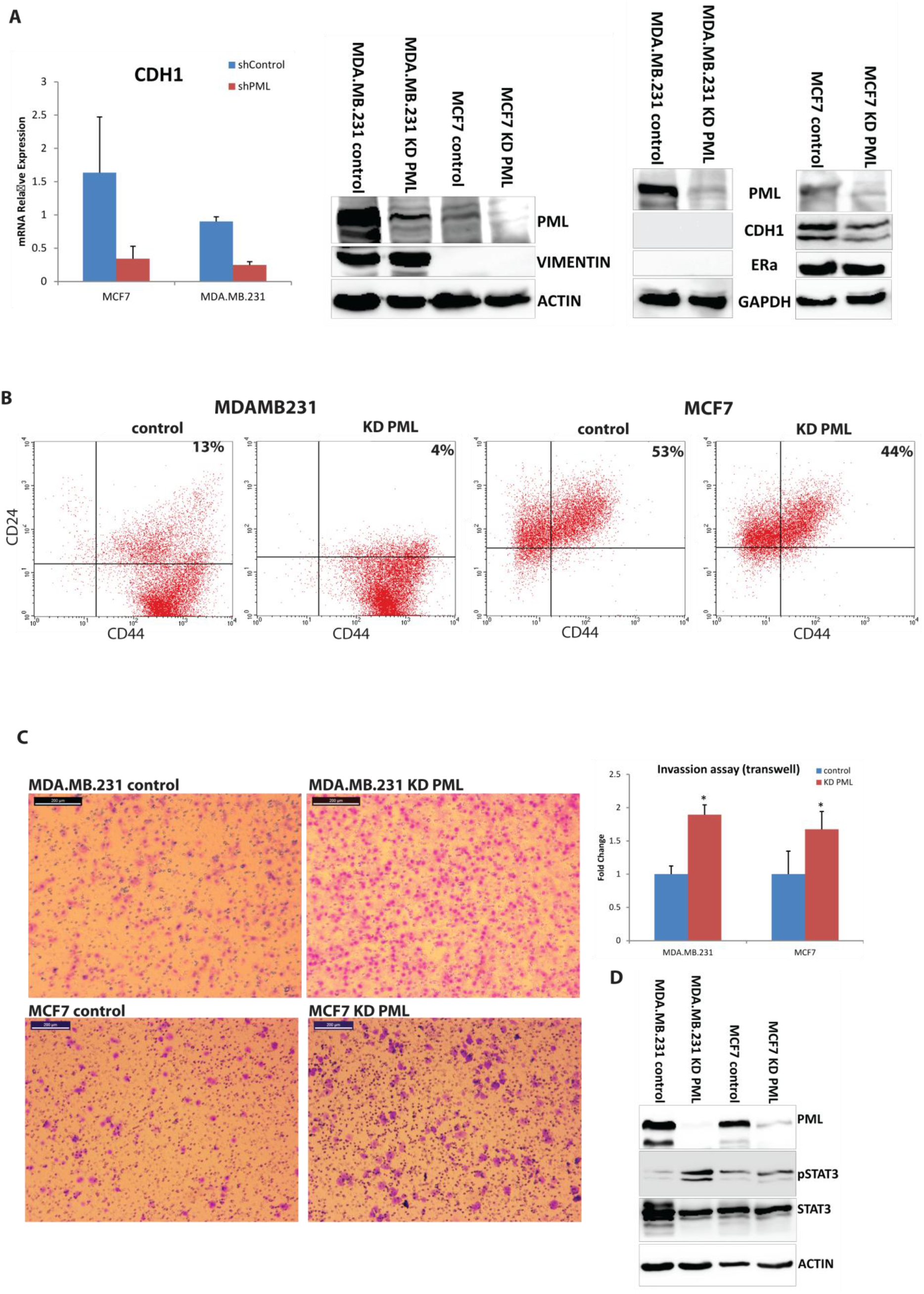
PML loss enhances mesenchymal properties in breast cancer cells. (A) qRT-PCR of MDA-MB-231 and MCF7, control and PML -KD cells of CDH1 expression (left). Results are mean ±SD from a triplicate experiment. Protein expression of a mesenchymal marker (Vimentin) (middle) and an epithelial (CDH1) (right). Actin and GAPDH are loading controls. (B) Flow cytometry for CD44 and CD24 surface expression in control and PML KD MDA-MB-231 (left panels) & MCF7 (right panels) cells. (C) Microscopy of migrated, MDA-MB-231 and MCF7 control or PML KD cells (Left panels). Quantitation (%) of crossing cells ±SD mean of one triplicate experiment (Right). t-test *p-value≤0.05. (D) Immunoblot for pSTAT3 protein (Tyr-705) and total STAT3.

**Figure 3.**
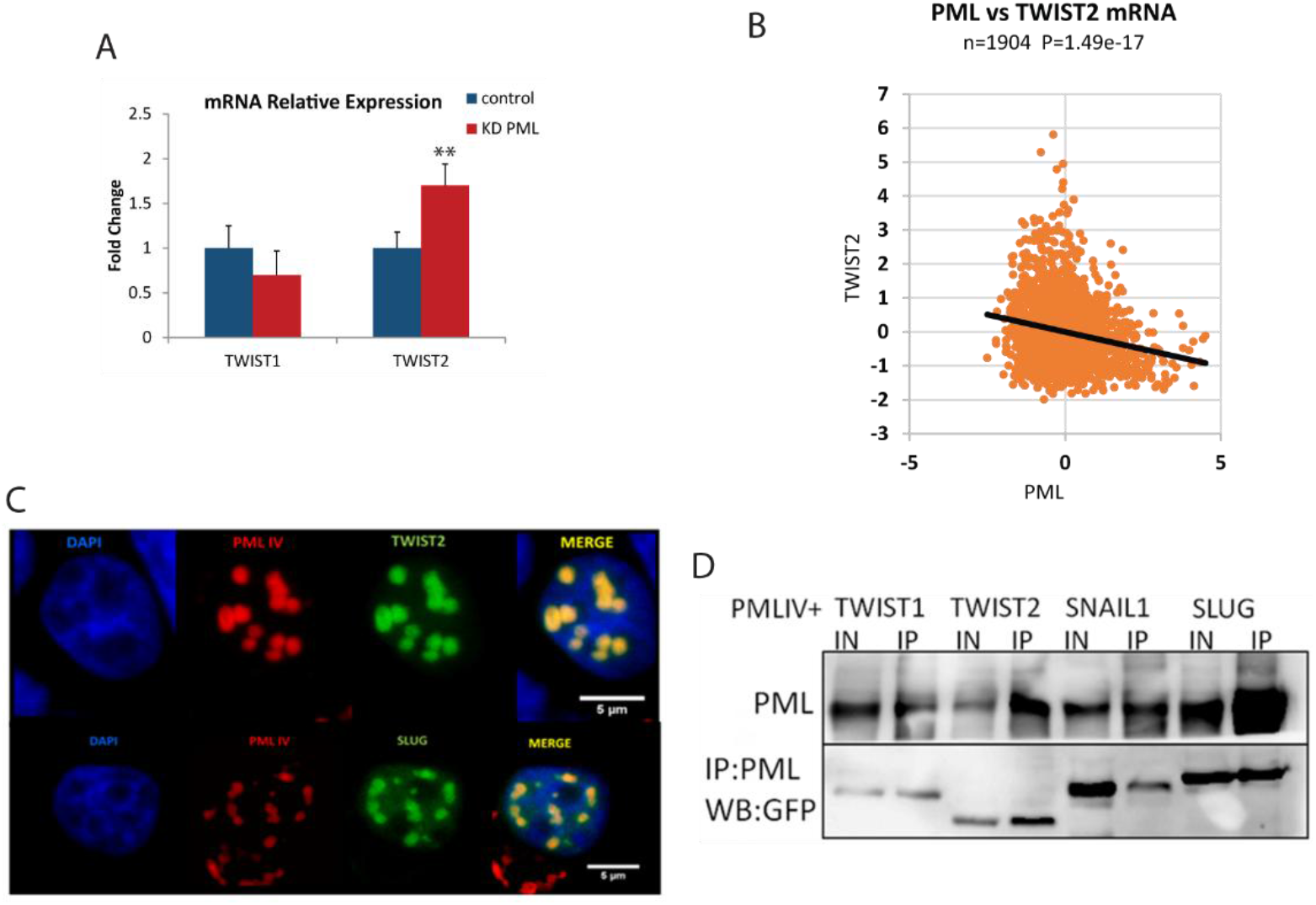
PML interacts with EMT factors and negatively correlates with TWIST2 expression. (A) qRT-PCR of MDA-MB-231, control and PML -KD cells for TWIST1 & TWIST2 expression. Results are mean ±SD from a triplicate experiment. t-test **p-value≤0.05. (B) Inverse correlation of PML –Twist2 RNA expression levels (log2 microarray from Metabric data set). (C) Confocal images of co-localization between PML isoform IV (fused to mRED) and TWIST2 or SLUG (fused to GFP). (D) Co-immunoprecipitation of PML with TWIST 1, TWIST2, SNAIL1 AND SLUG following co-expression in HEK293T cells. IN: input, IP: immunoprecipitated with a-PML antibody.

**Figure 4.**
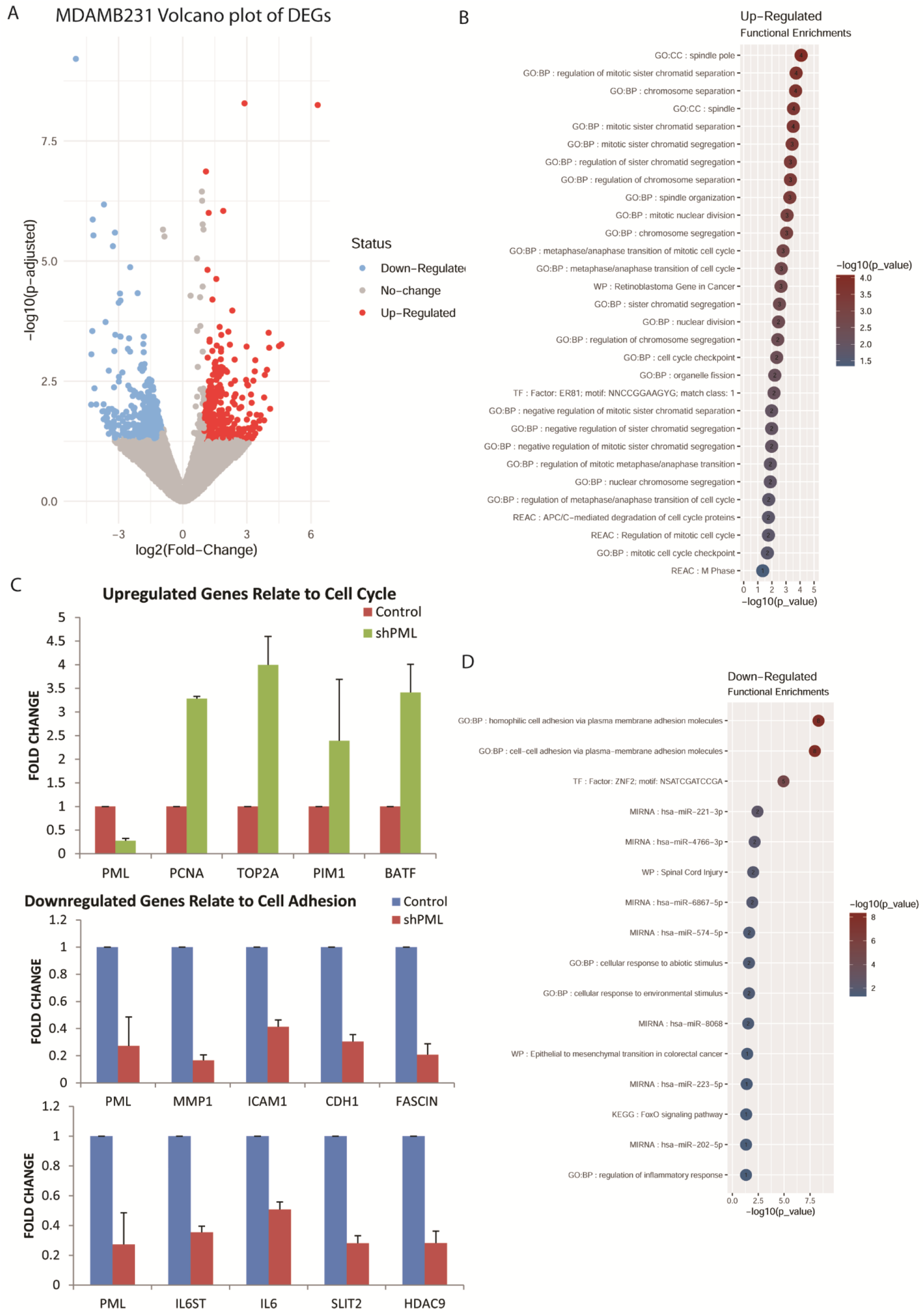
PML KD enriches expression of genes related to cell cycle and cell adhesion. (A) Volcano plot of differentially expressed genes in MDA-MB-231 KD PML, compared to control cells with p-value≤ 0.05, and log2 (FC)≥ 1. (B) Top enriched cellular processes of upregulated or down regulated (D) genes by PML loss in MDA-MB-231 cells. (C) Genes validated by qRT-PCR. Results are mean ±SD from a triplicate experiment.

**Figure 5.**
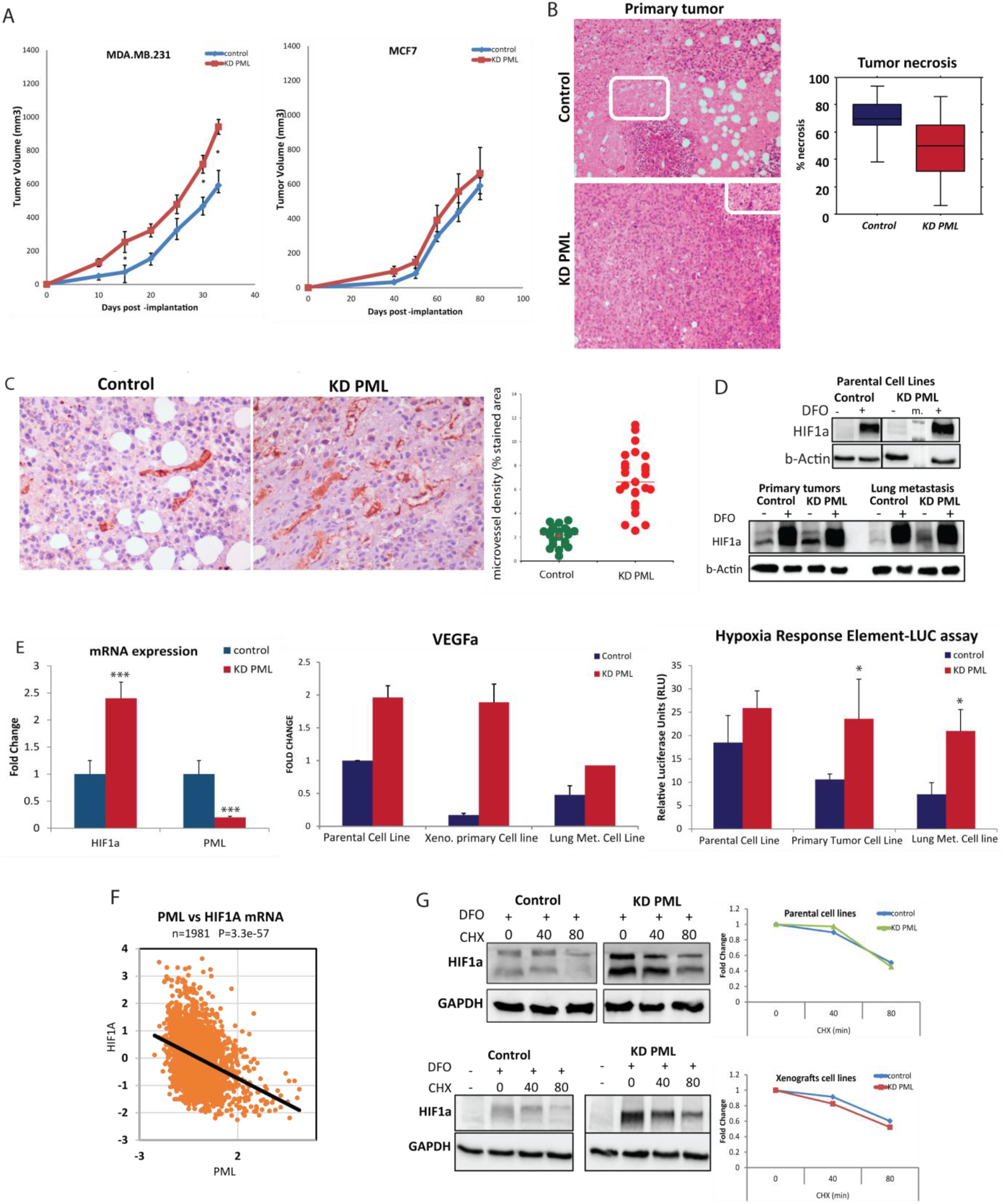
PML loss increased in vivo primary tumor growth and vascularization via HIF1a. (A) Tumor growth of MDA-MB-231 and MCF7, control (blue, n=7) & KD PML (red, n=7). Results show mean volume ±SD from one of two independent experiments with similar outcomes. *t-test p-value≤ 0.05. (B) Hematoxylin -Eosin at 100x magnification of primary MDA-MB-231 control & KD PML tumors (left). Rectangles indicate necrosis. Quantitation of primary tumor necrosis is presented by box plots of % necrotic area of primary tumors (n=7/group, Mann Whitney U,p-value ≤0.05) (right). (C) CD31 staining at 100x magnification of primary MDA-MB-231 control & KD PML tumors (left). Quantitation of primary tumor staining with CD31, indicative of angiogenesis (n=21 areas for control group and 25 areas for KD PML group/2-5 representative areas per tumor/5 tumors) (right), *t-test p-value ≤0.001). (D) Immunoblot for HIF1a protein in parental and xenograft derived lines from primary or lung metastatic MDA-MB-231 cells with or without DFO (Desferrioxamine 100 μM) treatment for 12h (E) Comparative expression of HIF1a gene in parental cell lines (left) and VEGF-α, a gene target of HIF1a, in parental or xenograft derived cell lines. Results are mean ±SD from a triplicate experiment (middle). HRE-LUC assay of control or PML KD parental cell lines, and primary tumor site or lung metastasis derived cell lines. Results are mean ±SD from one out of two triplicate experiments with similar outcomes (right). t-test *p-value≤0.05,***p-value≤0.01 (F) Inverse correlation of PML–HIF1a RNA expression levels (log2 microarray from Metabric data set). (G) Western blot of HIF1a in parental and xenograft derived lines from primary MDA-MB-231 cells treated with DFO for 4.5 h followed by treatment with cycloheximide (CHX). Graphs express HIF1a levels upon normalization for β-actin as fold over cells treated with CHX after DFO.

**Figure 6.**
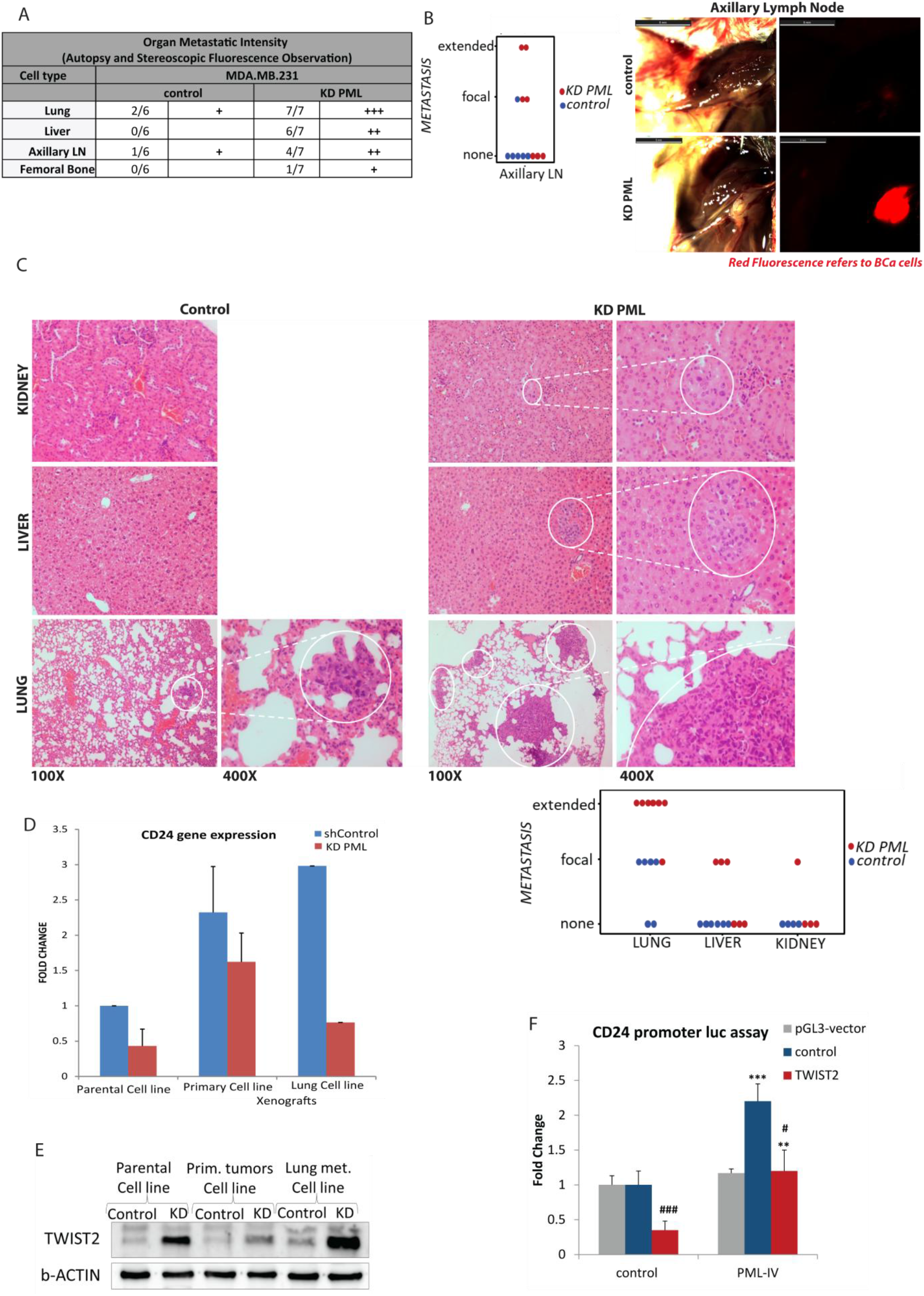
PML loss exacerbated the metastatic phenotype of MDA-MB-231 cells. (A) Summary table of metastases in mice with MDA-MB-231 control & MDA-MB-231 KD PML cells and grading by stereoscopic observation as +:positive, ++:focal and +++:extensive. (B) ImageJ quantification of axillary gland metastasis by fluorescence stereomicroscopy (χ2 test p-value ≤0.05). Representative axillary gland metastasis imaged by visible or fluorescence stereomicroscopy (right). (C) Top: Representative Kidney, Liver and Lung histologies of control and PML -KD engrafted mice. Hematoxylin-Eosin staining at 100x and 400x magnifications. Encircled areas indicate metastatic foci. Bottom: Quantitation and χ2 test of metastatic load in control and PML-KD engrafted mice for Lung (p=0.002), Liver (p=0.026) and Kidney (p=0.28). (D) CD24 mRNA expression in parental and xenograft derived cell lines. (E) TWIST2 protein expression in parental and xenograft derived cell lines. (F) CD24 promoter activity in control or TWIST2 cotransfected, untreated or dox –induced PML IV expressing cells. t-test ###p-value≤0.005, ***p-value≤0.005, #p-value≤0.05, **p-value≤0.05.

Thus, PML loss seems to intensify the existing mesenchymal phenotype while reducing the epithelial phenotype of MDA-MB-231 cells, whereas in MCF7 cells, leads to a decrease of the CDH1 epithelial marker without inducing mesenchymal marker expression. Because epithelial-mesenchymal transition (EMT) transcription factors (EMT-TFs) are mediators of the above phenotypic changes, we measured RNA levels of TWIST1 and TWIST2 key EMT factors and observed that PML loss caused a significant increase of TWIST2 expression at the RNA (Fig.3A) and protein level (Fig.6E). In line with this, the TCGA breast cancer (Metabric) data set [28] show a strong inverse correlation of RNA expression between PML and TWIST2 (Fig. 3B).

To further study the relationship of PML and EMT factors, PML-mCherry fusions and EMT factor-GFP fusions of TWIST1, TWIST2, SNAIL1 AND SLUG (SNAIL2) were co-expressed in HEK293T cells. PMLIV strongly co-localize with all EMT factors especially with TWIST2 and SLUG (Fig.3C), whereas PML III or PML I isoform did not (Suppl. Fig S1). A strong physical interaction of EMT factors and PML was further confirmed by co-immunoprecipitations (co-IPs). Figure 3D shows results from a representative co-IP using PMLIV. To define the region involved in the protein–protein interaction, we expressed TWIST2 truncations along with PMLIV followed by confocal microscopy and co-immunoprecipitation. Confocal results showed localization with the central region of TWIST2 that includes the bHLH. Indeed the HLH region alone (aa64 to aa124) showed a strong co-IP signal whereas the N- or C-terminal parts had minimal or no interaction ability (Suppl. Fig S2).

### 3.2. Changes in gene expression induced by PML ablation in breast cancer cells

To correlate the above changes in cell behaviour with potential effects of PML loss in gene expression, we performed transcriptomic analyses of control and PML KD cell lines. Specifically, silencing of PML deregulated about 600 genes at the 1.5 fold cut-off in MDA-MB-231 cells. Upregulated genes were enriched in cell cycle and cell division, whereas down-regulated genes showed enrichment in cell adhesion functions, Epithelial Mesenchymal Transition (EMT) and microRNAs that target tumour suppressors (Fig.4). A similar analysis of control and PML KD MCF7 cells identified 736 down- and 256 up-regulated genes. Upregulated genes were enriched in differentiation, morphogenesis, locomotion, and cell adhesion terms that involve NFKB1, RELA, and HIF1A factors while down regulated genes involve ECM organisation and cytokine signalling.Validation of deregulated expression for a number of those genes was done by qRT-PCR analysis (Suppl. Fig S3).

Comparisons of the two cell types showed little overlap in deregulated genes indicating a cellular context-specific response to PML loss. Common GO terms were related to stress response and developmental processes.

### 3.3. PML affects growth and hypoxic response of tumors derived from MDA-MB-231 xenografted cells

To study and correlate the above results with the *in vivo* tumor growth of PML-silenced cells, we performed engraftment in NSG mice. PML loss led to an increase of the tumour growth rate in MDA-MB-231 cells, but not MCF7 cells (Fig.5A). Histology of primary MDA-MB-231 KD PML tumour sites showed reduced necrosis relative to control group (Fig.5B). To test whether reduced necrosis might be due to enhanced vascularization we used anti CD31 antibody staining in sections of primary site tumors. PML KD tumors showed significantly higher microvessel density relative to the control group (Fig.5C). To further study these cells after their *in vivo* passage, we readapted tumor tissue back into cell culture and developed more than 16 lines from primary or metastastic sites. We examined the basal and (desferrioxamine) DFO induced HIF1A protein of parental and xeno-derived lines and found that PML KD had a 2 fold higher basal HI1A expression that was further intensified following *in vivo* growth (Fig.5D). Luciferase assays by transient transfection of the 5xHRE luc hypoxia response corroborated these results (Fig.5E right) PML loss resulted in a similar increase of HIF1A RNA and VEGF-α, a target of HIF1a and a main angiogenic factor, in both the parental and xeno-derived PML KD lines, (Fig.5E left). In agreement Inspection of clinical breast cancer data from TCGA–Metabric, showed a striking inverse correlation of PML and HIF1a (Fig.5F). Taken together these results suggest that PML loss facilitates HIF1A RNA expression to promote the acquisition of a hypoxic phenotype that is further enhanced in a hypoxic primary site relative to the lung metastatic growths. Because HIF1a is well known to be regulated at the protein stability level we next studied protein turn over in control and PML KD cells first induced with DFO and next with Cycloheximide (CHX). Similarly to basal HIF1A protein levels, DFO-mediated inhibition of degradation resulted in a proportional two fold increase in both parental and xeno-derived, either control or PML KD lines. A follow up inhibition of protein synthesis by cycloheximide showed that PML loss did not affect the protein degradation rate (about 80min half-life). Overall the above indicate that PML loss promotes Hypoxia response by mainly affecting both the RNA and protein expression but not turn over (Fig.5G).

### 3.4. The ablation of PML regulates the metastatic potential and organ preference of MDA-MB-231 cells

Gross observation and stereomicroscopy of autopsied tumor bearing mice showed that all (7/7) the PML KD expressing, MDA-MB-231 tumours, had a more intense metastatic load relative to controls in all the organs examined. More specifically all PML KD mice had many more lung metastatic foci, 6/7 were positive for liver metastases, 4/7 showed metastases in the axillary glands and one (1/7) showed metastasis to the bone marrow (Fig.6A) compared with tumors from control mice. These were further confirmed and quantified by stereomicroscopy and histology of axillary lymph node, liver, kidney and lung samples. Of note, lung was the most extensively affected site observed (Fig.6B,C) with an estimated 10 fold higher metastatic load in the PML KD (see also Fig.7A). MCF7 control or PML KD tumors showed no significantly different growth rate and no metastasis were found in either group (Suppl. Fig. S5). All PML control and KD lines derived from tumor tissue retained differential expression of PML RNA, and protein, similar to their parental cell lines (Suppl. Fig S4). Interestingly,MDA-MB-231PML KD cells or tumor –derived lines had increased TWIST2 protein expression (Fig.6E). In addition, they showed higher expression levels *of* recurrence-metastasis related markers, such as EpCAM[29] and CD49f [30] (Suppl. fig S4) and reduced epithelial CD24 RNA levels (Fig.6D).Using a transiently expressed CD24-Luc reporter we found that cotransfected TWIST2 reduced and expression of PML IV rescued luciferase activity (Fig.6F). Taken together, these results suggest that PML KD cells are enriched in mesenchymal properties consistent with a pro-metastatic phenotype because PML antagonises EMT factors by acting at both the RNA expression and protein function levels.

The increased lung metastatic ability of MDA-MB-231PML KD derived tumour lines (Fig.7A) was maintained upon secondary xenografting (Fig.7C). By qRT-PCR analysis of characteristic lung metastasis signature genes [14], namely ID1 and FSCN1, using the original and tumor derived cell lines we found that PML KD derived cells had higher expression than their parental cell lines (Fig.7B). Because results of tumor sphere formation (TSF) ability seemed not to be in line with *in vivo* tumor growth, we chose to examine in more detail the TSF ability of parental and tumor derived MDA–MB-231 cell lines under normoxic or hypoxic conditions. Results from these experiments show that passage one (P1) PML KD from either parental or tumor derived lines, had a lower TSF ability (20-30%) than controls, under either normoxic or hypoxic (3% oxygen) conditions, although in the latter a general increase of TSF was found. However, during subsequent passages the TSF ability of PML KD cells increased whereas that of the control groups decreased. At P3, parental, primary tumor and lung derived PML KD cells exceeded their controls in pair wise comparisons (Fig.7D).

**Figure 7.**
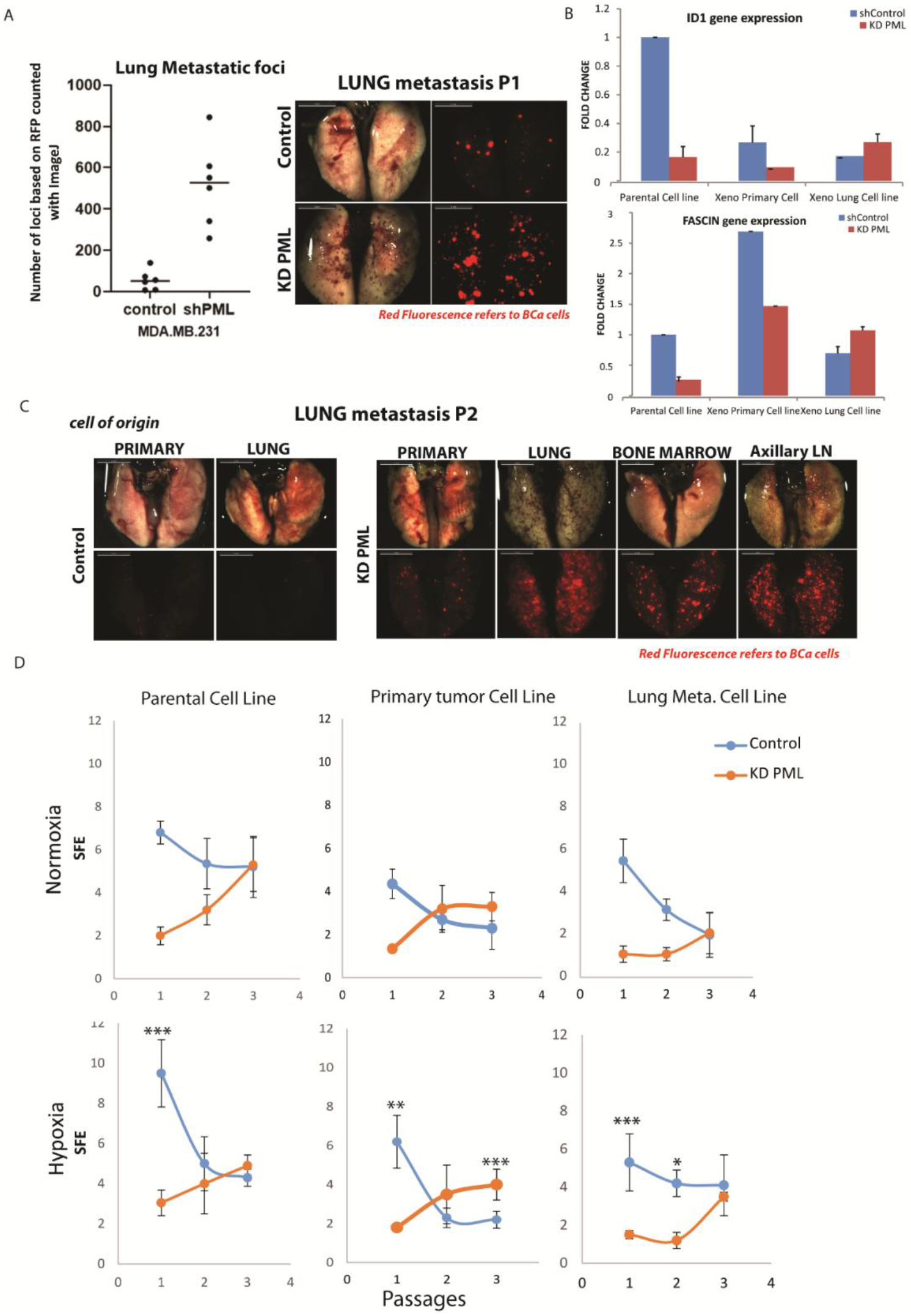
PML loss promotes lung organotropism and sustains long-term TSF ability. (A) Assessment of Lung metastatic load of control and PML –KD engrafted mice by comparing (t-test, p-value≤0.05) total foci counts (left) and representative stereo microscopy image (middle). (B) Comparative expression of lung “metastasis signature”[14] (LMS) genes ID1 and FSCN1 in control and KD PML, parental and xenografts derived cell lines. Results are mean ±SD from a triplicate experiment. (C) Lung stereo microscopy from second passage (P2) of lung metastases following re-injection of first passage derived cells. (D) Percentage (%) of cells forming spheres (SFE), when cultured under normoxic or hypoxic conditions from parental and xenograft derived cell lines through sequential passaging P1, P2, P3. Results show mean SFE ±SD from one out of at least two experiments in quadruplicate. t-test *p-value≤0.05,**p-value≤0.01, ***p-value≤0.0001.

## 4. DISCUSSION

To investigate the role of PML in breast cancer we produced Knock Down lines of the claudin-low MDA-MB-231 and the luminal-A MCF7 and studied their physiology and transcriptomic profiles.

Our previous studies have shown that induced overexpression of PMLIV in MDA-MB-231 cells strongly inhibited cell proliferation and tumor sphere formation (TSF) ability[11]. While PML silencing of all PML isoforms in those cell types caused a slight increase of the cell growth rate in monolayer culture, yet their tumor sphere ability was compromised (25-30% of the control line) implying a role for PML in their maintenance of stem cell like properties. In addition, PML loss changed the MCF7 tumor spheres from their known smooth spherical morphology[24] to a more grape one, reflecting a change in the expression of cell–cell interacting proteins. The ablation of PML caused a shift in both cell line towards a more mesenchymal appearance that has been previously linked to increased cell invasiveness[31,32] and tumor recurrence or metastasis[33,34]. In accordance with this, PML deficient cells showed increased migration capacities. The increase in migration activity was more pronounced in MDA MB 231 cells and correlated with higher phosphorylated Stat3[35].

To correlate these changes with RNA expression profiles, we report here for the first time a transcriptomic analysis upon silencing of PML. The transcriptomic analysis reflected the differences in the physiology of these cell types. The ablation of PML in MDA-MB-231 cells caused the repression of genes related to cell adhesion and ECM. On the other hand, genes related to cell cycle, mitosis and cell division were upregulated. A similar analysis in MCF7 cells identified developmental–differentiation related genes to be upregulated and cytokine and extracellular matrix related genes to be repressed upon PML KD. Interestingly, earlier studies of transient PML KD in HUVEC cells identified some common GO enriched classes such as cell cycle and locomotion[36].

Together the phenotype and deregulated gene expression profile suggest that PML deficient MDA-MB-231 cells are expected to be more aggressive than their parental cells. This contrasts with the reduction in TSF that is thought to measure of tumor initiating cells (TICs) or cancer stem cell activity. To clarify this we xenografted control and PML KD cells in NSG mice and monitored their growth in the primary site and development of metastasis. PML KD did not affect significantly the local tumor growth or the non/low metastatic properties of MCF7 cells. However, MDA-MB-231 KD PML cells developed tumors faster when compared with the controls. This observation contradicts the *in vitro* TSF assay suggesting that the growth defect in PML loss can be reversed during *in vivo* tumor growth. Indeed, upon sequential tumor sphere passaging, PML KD cells showed a gradual increase of their TSF ability concomitant with a decrease of the control TSF. These results show that PML KD cells retain higher long-term TSF ability especially when taken from primary tumor sites and/or hypoxic conditions in agreement with their higher *in vivo* tumor growth rate and metastasis. Taken together these results suggest that PML KD cells acquire higher TIC properties *in vivo* and show a more aggressive phenotype. Tissue hypoxia seems to further intensify this process. These results partly disagree, with the observations of Ponente et al.[37], Martin-Martin et al.[38] And Arreal et al.[39] that describe a pro-oncogenic role of PML in breast cancer. Although presently undetermined these differences may be caused by distinct MDA-MB-231 subline differences or the effect of long term and strong PML silencing in our lines.

EMT factors is a large family of proteins with important roles in normal development often found to be deregulated in cancer[40] in response to upstream activators[41]. The role of interaction of PML with EMT factors seems to be complex. Cytoplasmic PML promotes EMT via potentiation of TGF b signaling[42,43]. We described here for the first time that both bHLH (TWIST) and Zn-finger (SNAIL) type EMT factors bind and co-localize to PML IV and to a lesser extent to PML I and III driven nuclear bodies. Interaction with TWIST2 requires the presence of the bHLH domain and may result in altered dimerization or DNA binding in a way similar to the inhibitory action of IDs on EMT actors. Functionally PML overexpression reverses the inhibition of TWIST2 on the CD24 promoter. In addition, PML KD leads to increased TWIST2 RNA and protein expression in parental tumors and the lung metastases in line with a strong inverse correlation of PML and TWIST2 expression in a large breast data set (TCGA-Metabric).

An important observation of this study was the increased metastatic properties of PML deficient MDA-MB-231 cells to various sites and a stronger preference for the development of lung metastasis. Earlier studies have identified a group of genes in MDA-MB-231 cells expressed at higher levels in cell lines derived from lung metastases (lung metastasis signature, LMS) relative to their parental cell line[14]. The differentially expressed PML KD genes and the LMS list (shown in suppl. Table S1) contain 11 common genes (Shown in Suppl. Table S2). Interestingly FSCN1 and ID1 showed a change in their expression pattern resembling that of LMS. The ID gene family has important roles in development and their deregulated expression is connected to poor prognosis. Increased expression of ID1 has been associated with both primary tumor formation and lung colonization[44] presumably via mesenchymal -epithelial transition essential for the latter process[45]. Earlier studies demonstrated that MDA-MB-231 are genetically heterogeneous and *in vitro* selection for more aggressive populations with altered genome structure have been reported[46]. Similarly, sequential *in vivo* growth of MDA-MB-231 selects for highly metastatic sublines enriched in pre-existing or newly arising genetic variants such as BRAF and KRAS[47] as well as ALPK2 and RYR1[48] that correlate with high organ specific metastatic ability. Although not directly demonstrated it is likely that PML deficiency accelerates this *in vivo* selection process. In the present studies, PML affected both the primary and metastatic tumor growth. Searching for metastatic suppressor genes (MSG) we found that the ablation of PML resulted in the up - or down regulation of a small number of MSGs[49]. but their functional significance remains to be determined.

Tumors produced by MDA-MB-231 PML KD cells showed lower necrotic content suggesting higher vascular supply. In agreement with this, CD31 immunocytochemistry showed significant increase of microvessel density in PML KD primary tumors. In agreement with this, parental- or lines generated by culturing PML KD tumors expressed higher levels of HIF1a protein and its target, VEGFα that may account for a higher tumor angiogenic activity. We show here that PML loss enhanced HIF1A and consequently VEGF RNA expression without affecting the protein turn over unlike its role in promoting HIF1A protein synthesis in a mouse KO model. [50]. Because HIF1A is mainly regulated at the protein level understanding of its transcriptional regulation is less well studied. Nevertheless there is evidence of various regulatory pathways that can activate HIF transcription[51] including STAT3[52]. Strikingly, we have observed STAT3 activation in the PML KD -MDA MB231 cells. PML is a known IFN inducible gene[53]. Moreover STAT3 KD down-regulates PML[38]. Conversely, PML blocks STAT3 via direct binding [54] and blocking its IL6-mediated activation[55]. In addition PML loss prolongs nuclear STAT3 life[56]. Thus PML stands in the middle of a negative feedback n network with cytokines/IFNs and STAT3. Importantly STAT3 regulates both EMT and HIF1A [52] and may also activate angiogenic target genes either acting alone or in a network with HIF1a[57]. Among others, STAT3 induces TWIST2 expression[58]. These results suggest that STAT3 may be the common culprit that promotes both EMT/migration and HIF/angiogenesis to generate a higher metastastic load. Unrestrained STAT3 activity in PML deficiency facilitates both EMT and angiogenic signaling for in PML enhanced aggression and metastasis of TNBC cells.

## 5. Conclusions

Collectively PML impedes tumorigenicity by regulating the mesenchymal-invasive and angiogenic properties and the metastatic tropism of MDA-MB-231 cells. In addition PML may control the long term survival of TICs. Finally, PML loss may accelerate *in vivo* evolution of genetically heterogeneous population to acquire a more aggressive-metastatic phenotype. Mechanistically this is linked to the long term maintenance of tumor initiating cells, potentiation of their mesenchymal phenotype and a hypoxia-angiogenic switch. Hence, PML expression may serve as a valuable cancer biomarker.

## Supporting information

new suppl figures

new suppl tables

primers and antibodies used in this paper

## Abbrevations

PML: promyelocytic leukemia protein
EMT: epithelial-mesenchymal transition
HIF1a: Hypoxia-inducible factor 1-alpha

## Data Availability Statement

RNA-seq datasets have been deposited in SRA database under accession number PRJNA826854.

## Author contributions

AV designed and performed experiments, data analysis, discussion and writing; SS, NS and TM contributed to the experimental work and discussion; ED performed immunohistochemistry in paraffin sections and histological analysis; CN performed bioinformatic analysis; AK contributed to discussion and writing; JP supervised the study and contributed to writing.

## Acknowledgements

We would like to thank E.Tachmatzidi and O.Galanopoulou for helping with the *in vivo* experiments. We would also like to thank G.Vretzos and D.Tsoukatou for technical assistance, as well as the Genomic Facility of IMBB for the gene expression profiling, the Confocal Microscopy Facility and High Content Screening Facility of IMBB for imaging and data analysis respectively. We also thank M.Dimakopoulou, I.Charatsidou and C.Fountas for helping with this work.

## Conflict of interest

The authors declare no conflicts of interest.

## Funding

The authors of this work were funded by “DINNESMIN-MIS 5032840”, “EDBM103-MIS 5048184” and IMBB-FORTH internal funding.

## Notes

### Competing Interest Statement

The authors have declared no competing interest.

### Summary of Updates

We provide new experiments that further support our conclusions and provide the mechanism of PML actions: a. PML ablation increases TWIST2 and HIF1a RNA and protein levels. These results are corroborated by showing an inverse correlation of PML and TWIST2 or HIF1a RNA levels in a large cohort of more than 1900 breast cancer patients from the TCGA data base b. PML binds to TWIST2 and in addition rescues its suppressor action on the CD24 gene promoter. Thus PML antagonises TWIST2 both at the RNA expression and protein action levels. c Because HIF1a is well known to be regulated by protein degradation we investigated this and found that PML loss, does not affect protein turn over as suggested by other studies in mouse models.

