## Supplementary material for "Promyelocytic Leukemia Protein regulates Angiogenesis and Epithelial-Mesenchymal Transition to limit metastasis in MDA-MB-231 breast cancer cells": new suppl figures

**Figure S1 (related to figure 3):** Immunoprecipitation of PML protein in HEK293T cells, following overexpression of PML I & III isoforms with TWIST2-GFP western blot analysis. Colocalization of PMLI & III and TWIST2 under a confocal microscope, PMLI & III (red), TWIST2 (green).

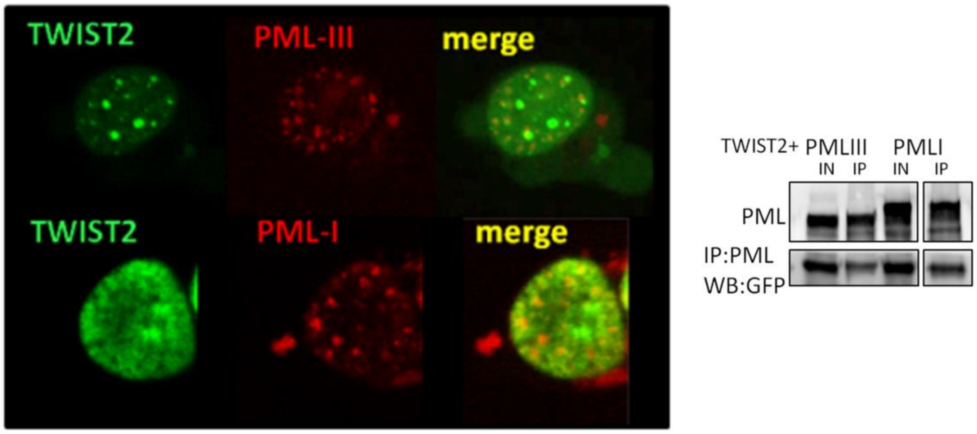


**Figure S2 (related to figure 3):** PML protein immunoprecipitation experiment in HEK293T cells following overexpression of PMLIV and TWIST2 fragments; Western blot analysis for TWIST2-GFP; Colocalization of PMLIV and TWIST2 parts under a confocal microscope, PMLIV (red), TWIST2 (green); GFP-NLS (green-negative control)

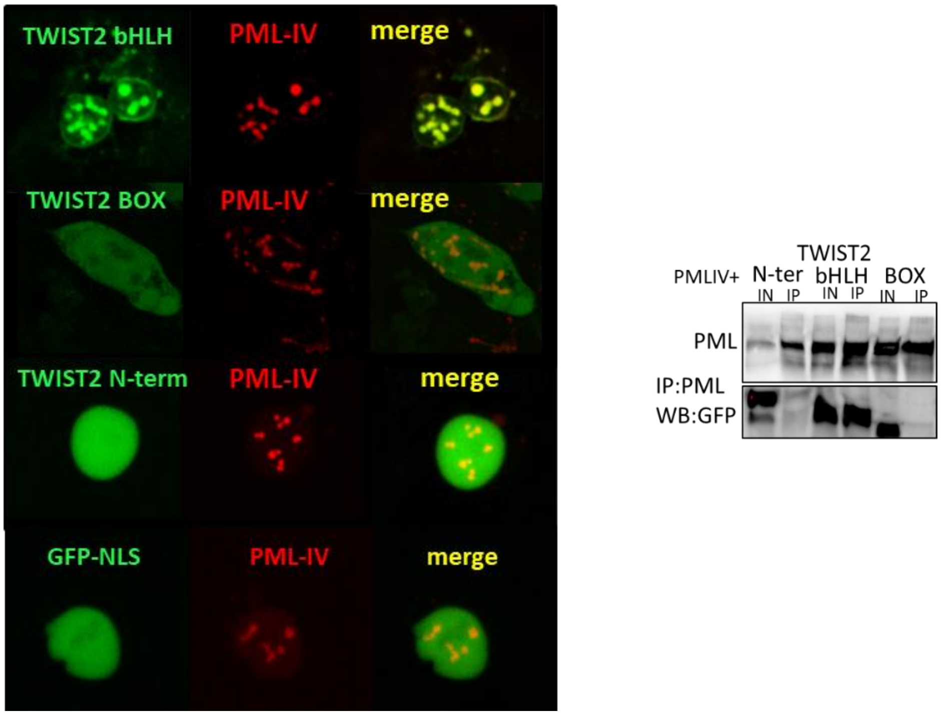

**<u>Figure S3 (related to figure 2):</u>** Bioinformatics analysis of MCF7.

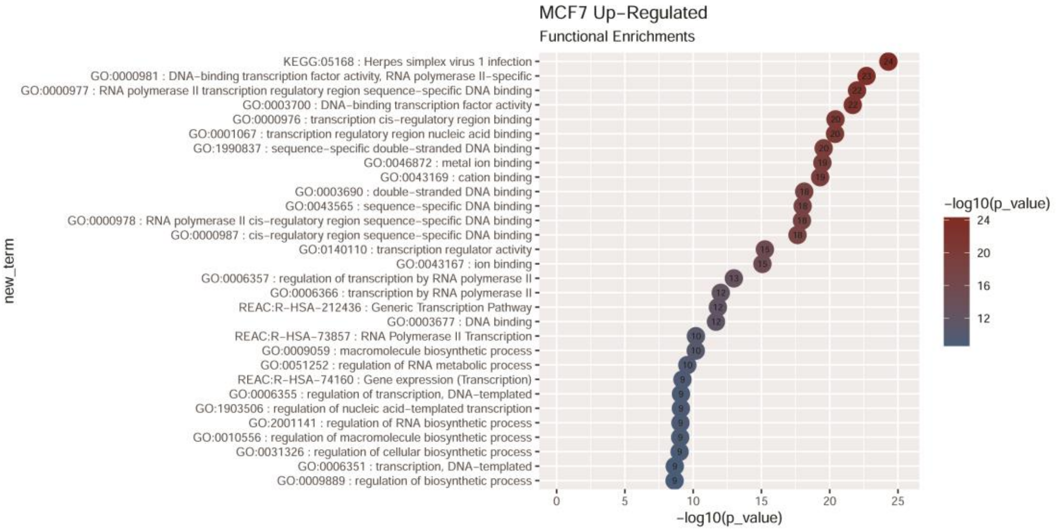


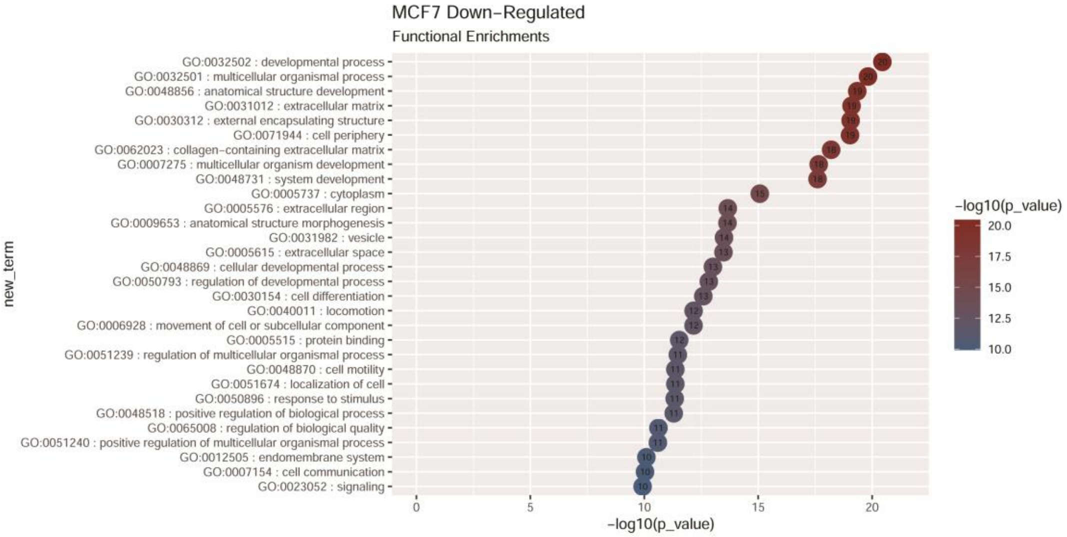


<u>MCF7 gene expression validations</u>

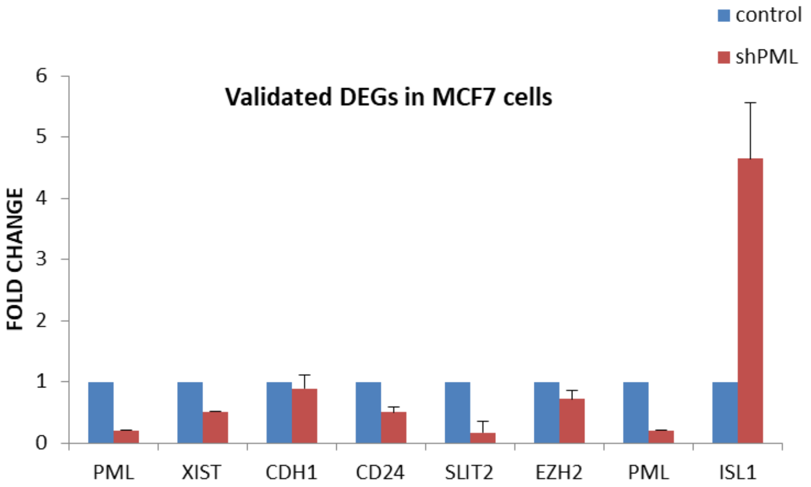

**Figure S4 (related to figure 6):** (A) Expression of PML gene and protein (qRT-PCR/Western blot/immunohistochemical analysis) in the experimental study groups MDA.MB.231 & MDA.MB. KD PML. (B) PML protein expression for primary tumors and their metastases. (C)Expression of EpCAM and CD49f integrin (ITGA6) in cell lines and primary tumors compared with control groups.

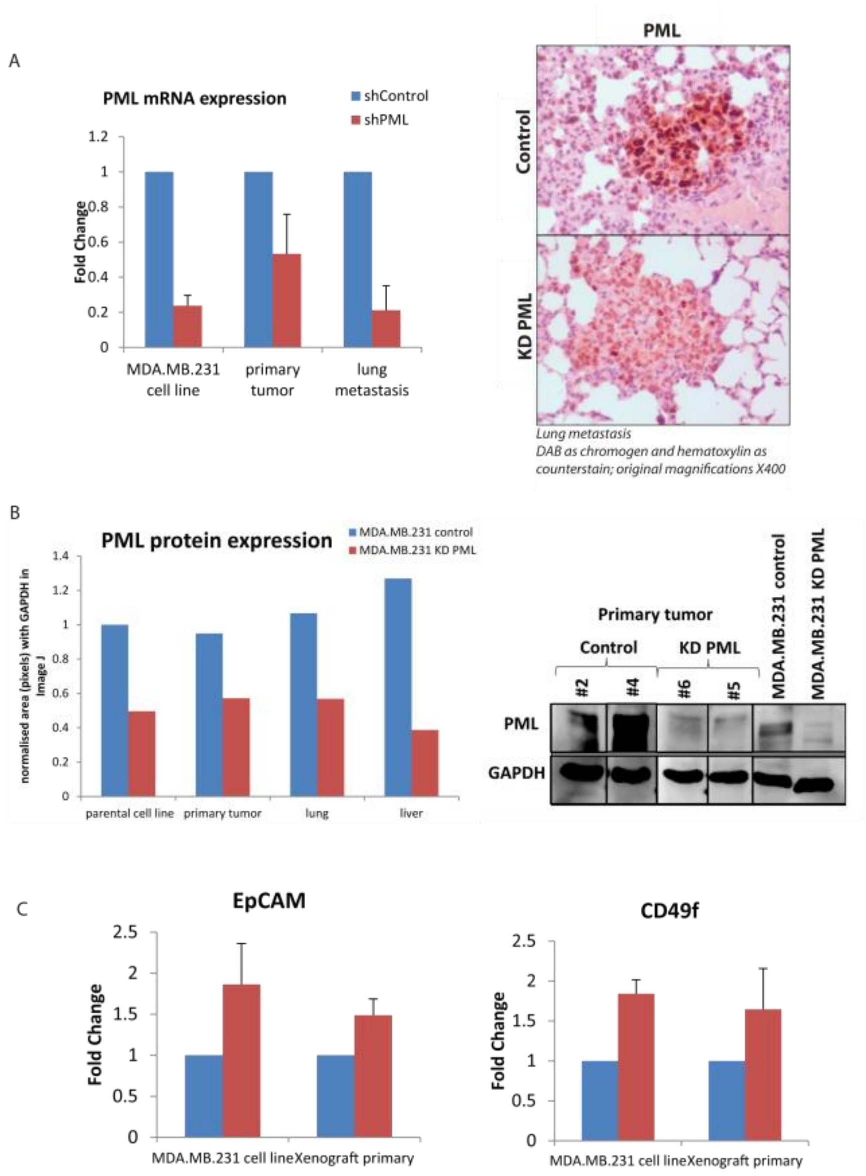

**Figure S5 (relatedto figure 6):** Lungs from the two study groups MCF7 & MCF7 KD PML. The observation was made on a fluorescence stereoscope (A). Summary table with the metastases that the experimental animals showed in the study groups (B). Expression of PML protein in animals indicative of isolated protein material from primary tumors (C).

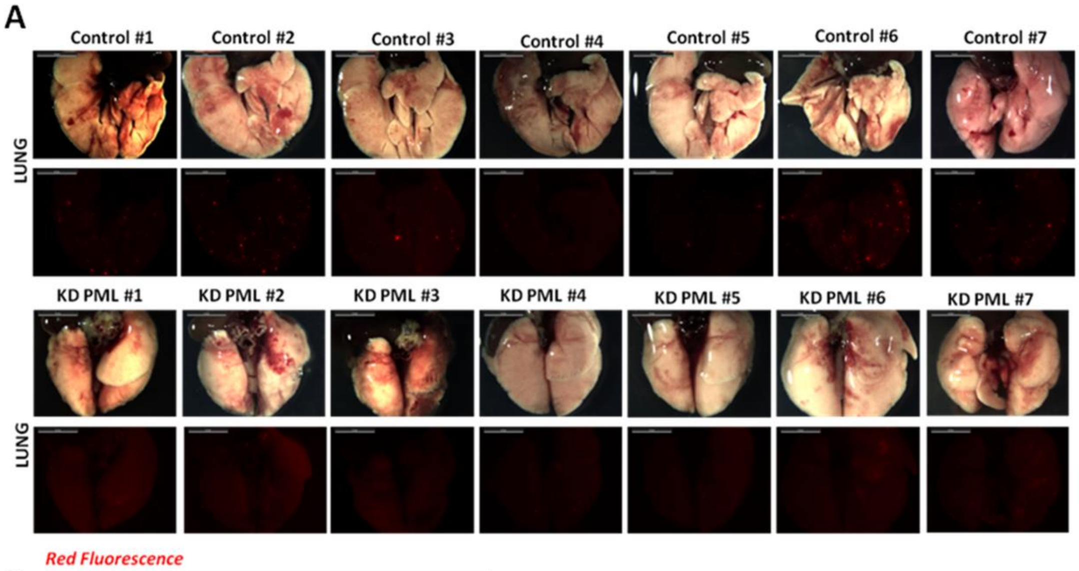


B

| Organ Metastatic Intensity (Autopsy and Stereoscopic Fluorescence Observation) | | | | |
|---|---|---|---|---|
| Cell type | MCF7 | | | |
| | control | | KD PML | |
| Lung | 7/7 | ++ | 7/7 | + |
| Liver | 1/7 | + | 2/7 | + |
| Axillary LN | 0/7 | | 1/7 | + |
| Femoral Bone | 0/7 | | 0/7 | |

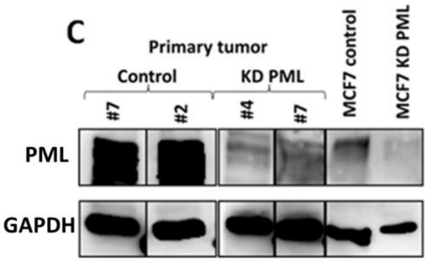
