## Supplementary material for "Promyelocytic Leukemia Protein regulates Angiogenesis and Epithelial-Mesenchymal Transition to limit metastasis in MDA-MB-231 breast cancer cells": new suppl tables

**Supplementary Tables**

**Table S1 (related to figure 7): List of 95 genes expressed in pulmonary metastases from MDA.MB.231 sequential cells (P2-LM2) identified in patients with primary tumors expressing or not these genes (Rosetta training set). *p-value ≤0.05.*[14] compared with parental cell line MDA-MB-231 differentially expressed genes (DEGs).**

| Fold Change | Gene Title | Gene Symbol |
|---|---|---|
| 407.01 | secreted protein, acidic, cysteine-rich (osteonectin) | SPARC |
| 147.27 | protein tyrosine phosphatase, receptor type, N polypeptide 2 | PTPRN2 |
| 97.07 | protein tyrosine phosphatase, receptor type, N polypeptide 2 | PTPRN2 |
| 58.71 | colony stimulating factor 3 (granulocyte) | CSF3 |
| 48.52 | interleukin 13 receptor, alpha 2 | IL13RA2 |
| 33.05 | killer cell lectin-like receptor subfamily C, member 1 /// killer cell lectin-like receptor subfamily C, member 2 | KLRC1 /// KLRC2 |
| 20.03 | collagen, type I, alpha 1 | COL1A1 |
| 15.67 | protein tyrosine phosphatase, receptor type, N polypeptide 2 | PTPRN2 |
| 14.65 | melanoma antigen, family D, 4 /// melanoma antigen, family D, 4 | MAGED4 |
| 13.50 | glycophorin C (Gerbich blood group) | GYPC |
| 13.35 | matrix metalloproteinase 1 (interstitial collagenase) | MMP1 |
| 12.82 | kynureninase (L-kynurenine hydrolase) | KYNU |
| 8.99 | Epiregulin | EREG |
| 7.43 | tenascin C (hexabrachion) | TNC |
| 6.77 | Interferon stimulated gene 20kDa | ISG20 |
| 6.75 | Aldehyde dehydrogenase 3 family, memberA1 | ALDH3A1 |
| 6.35 | collagen, type VI, alpha 1 | COL6A1 |
| 6.34 | keratin, hair, basic, 1 | KRTHB1 |
| 6.29 | kynureninase (L-kynurenine hydrolase) | KYNU |
| 6.23 | prostaglandin-endoperoxide synthase 2 (prostaglandin G/H synthase and cyclooxygenase) | PTGS2 |
| 5.83 | Lysosomal-associated multispanning membrane protein-5 | LAPTM5 |
| 5.74 | chromosome 10 open reading frame 116, adipose specific 2 | C10ORF116 |
| 5.29 | apolipoprotein B mRNA editing enzyme, catalytic polypeptide-like 3G | APOBEC3G |
| 5.02 | platelet-derived growth factor alpha polypeptide | PDGFA |
| 4.86 | roundabout, axon guidance receptor, homolog 1 (Drosophila) | ROBO1 |
| 4.63 | serine (or cysteine) proteinase inhibitor, clade E (nexin, plasminogen activator inhibitor type 1), member 2 | SERPINE2 |
| 4.56 | SPANX family, member C | SPANXC |
| 4.56 | angiopoietin-like 4 | ANGPTL4 |
| 4.55 | fascin homolog 1, actin-bundling protein (Strongylocentrotus purpuratus) | FSCN1 |
| 4.47 | jagged 1 (Alagille syndrome) | JAG1 |
| 4.45 | SRY (sex determining region Y)-box 4 | SOX4 |
| 4.40 | SPANX family, member B1 /// SPANX family, member C | SPANXB1 /// SPANXC |
| 4.26 | Rho GDP dissociation inhibitor (GDI) beta | ARHGDIB |
| 4.24 | collagen, type VI, alpha 1 | COL6A1 |
| 4.21 | SPANX family, member B1 | SPANXB1 |
| 4.16 | Interferon stimulated gene 20kDa | ISG20 |

| | | |
|---|---|---|
| 4.01 | glutaminyl-peptide cyclotransferase (glutaminyl cyclase) | QPCT |
| 3.99 | fascin homolog 1, actin-bundling protein (Strongylocentrotus purpuratus) | FSCN1 |
| 3.89 | chemokine (C-X-C motif) ligand 1 (melanoma growth stimulating activity, alpha) | CXCL1 |
| 3.85 | matrix metalloproteinase 2 (gelatinase A, 72kDa gelatinase, 72kDa type IV collagenase) | MMP2 |
| 3.76 | doublecortin and CaM kinase-like 1 | DCAMKL1 |
| 3.71 | Stomatin | STOM |
| 3.62 | G protein-coupled receptor 153 | GPR153 |
| 3.59 | mannosidase, alpha, class 1A, member 1 | MAN1A1 |
| 3.57 | chromosome 14 open reading frame 139 | C14orf139 |
| 3.54 | caspase 1, apoptosis-related cysteine protease (interleukin 1, beta, convertase) | CASP1 |
| 3.42 | immunoglobulin superfamily, member 4 | IGSF4 |
| 3.41 | latent transforming growth factor beta binding protein 1 | LTBP1 |
| 3.24 | kynureninase (L-kynurenine hydrolase) | KYNU |
| 3.24 | nuclear receptor subfamily 2, group F, member 1 | NR2F1 |
| 3.21 | epithelial membrane protein 1 | EMP1 |
| 3.21 | Lysosomal-associated multispanning membrane protein-5 | LAPTM5 |
| 3.17 | solute carrier family 22 (organic cation transporter), member 1-like antisense | SLC22A1LS |
| 3.15 | epithelial membrane protein 1 | EMP1 |
| 3.12 | microfibrillar-associated protein 2 | MFAP2 |
| 3.10 | inhibitor of DNA binding 1, dominant negative helix-loop-helix protein | ID1 |
| 3.10 | solute carrier organic anion transporter family, member 4A1 | SLCO4A1 |
| 3.07 | CCR4-NOT transcription complex, subunit 2 | CNOT2 |
| 3.07 | Activating transcription factor 1 | ATF1 |
| 3.06 | zinc finger protein 185 (LIM domain) | ZNF185 |
| 3.02 | hypothetical protein LOC221810 | LOC221810 |
| 0.33 | ATPase, Class VI, type 11A | ATP11A |
| 0.33 | muscleblind-like 2 (Drosophila) | MBNL2 |
| 0.33 | isocitrate dehydrogenase 2 (NADP+), mitochondrial | IDH2 |
| 0.33 | olfactomedin-like 2A | OLFML2A |
| 0.32 | neural precursor cell expressed, developmentally down-regulated 9 | NEDD9 |
| 0.32 | cofactor required for Sp1 transcriptional activation, subunit 2, 150kDa | CRSP2 |
| 0.32 | colony stimulating factor 2 receptor, alpha, low-affinity (granulocyte-macrophage) | CSF2RA |
| 0.32 | likely ortholog of mouse limb-bud and heart gene /// likely ortholog of mouse limb-bud and heart gene | LBH |
| 0.31 | molybdenum cofactor sulfurase | MOCOS |
| 0.31 | major histocompatibility complex, class II, DP alpha 1 | HLA-DPA1 |
| 0.30 | nicotinamide N-methyltransferase | NNMT |
| 0.30 | membrane-bound transcription factor protease, site 2 | MBTPS2 |
| 0.30 | claudin 4 | CLDN4 |
| 0.30 | EGF-containing fibulin-like extracellular matrix protein 1 | EFEMP1 |
| 0.30 | epoxide hydrolase 1, microsomal (xenobiotic) | EPHX1 |
| 0.30 | tumor necrosis factor (ligand) superfamily, member 10 | TNFSF10 |
| 0.29 | muscleblind-like 2 (Drosophila) | MBNL2 |
| 0.29 | TBC1 domain family, member 4 | TBC1D4 |
| 0.28 | myosin, heavy polypeptide 10, non-muscle | MYH10 |

| | | |
|---|---|---|
| 0.27 | receptor tyrosine kinase-like orphan receptor 1 | ROR1 |
| 0.27 | dishevelled associated activator of morphogenesis 1 | DAAM1 |
| 0.26 | haloacid dehalogenase-like hydrolase domain containing 1A | HDHD1A |
| 0.25 | glutathione S-transferase M4 | GSTM4 |
| 0.25 | LOC388483 | --- |
| 0.24 | gelsolin (amyloidosis, Finnish type) | GSN |
| 0.24 | myosin, heavy polypeptide 10, non-muscle | MYH10 |
| 0.24 | EGF-like repeats and discoidin I-like domains 3 | EDIL3 |
| 0.23 | major histocompatibility complex, class II, DP beta 1 | HLA-DPB1 |
| 0.23 | major histocompatibility complex, class II, DR beta 3 | HLA-DRB3 |
| 0.23 | major histocompatibility complex, class II, DR beta 3 | HLA-DRB3 |
| 0.23 | aryl-hydrocarbon receptor nuclear translocator 2 | ARNT2 |
| 0.22 | nerve growth factor, beta polypeptide | NGFB |
| 0.21 | retinoic acid receptor responder (tazarotene induced) 3 | RARRES3 |
| 0.21 | nicotinamide N-methyltransferase | NNMT |
| 0.21 | EGF-containing fibulin-like extracellular matrix protein 1 | EFEMP1 |
| 0.18 | calcium/calmodulin-dependent serine protein kinase (MAGUK family) | CASK |
| 0.18 | Major histocompatibility complex, class II, DP alpha 1 | --- |
| 0.17 | matrilin 2 | MATN2 |
| 0.16 | par-6 partitioning defective 6 homolog beta (C. elegans) /// par-6 partitioning defective 6 homolog beta (C. elegans) | PARD6B |
| 0.16 | serine protease inhibitor, Kazal type 4 | SPINK4 |
| 0.16 | colony stimulating factor 1 (macrophage) | CSF1 |
| 0.16 | complement component 4 binding protein, beta | C4BPB |
| 0.14 | lymphocyte antigen 6 complex, locus E | LY6E |
| 0.13 | major histocompatibility complex, class II, DP alpha 1 | HLA-DPA1 |
| 0.12 | chromosome 6 open reading frame 108 | C6orf108 |
| 0.10 | acetylserotonin O-methyltransferase-like | ASMTL |
| 0.09 | ATP-binding cassette, sub-family C (CFTR/MRP), member 3 | ABCC3 |
| 0.08 | chemokine (C-X-C motif) receptor 4 | CXCR4 |
| 0.07 | cystatin F (leukocystatin) | CST7 |
| 0.07 | KIAA1199 | KIAA1199 |
| 0.06 | chemokine (C-X-C motif) receptor 4 | CXCR4 |
| 0.04 | par-6 partitioning defective 6 homolog beta (C. elegans) | PARD6B |

**Table S2 (related to figure 7): List of 15 common genes between Lung signature gene set [14] and MDA-MB-231 DEGs.**

| Gene name | Fold Change in MDAMB231 parental cell line | Fold change in Minn et al 2005 |
|---|---|---|
| C4BPB | 0.460635 | 0.16 |
| CASP1 | 4.5 | 3.54 |
| EDIL3 | 2.25228 | 0.24 |
| EMP1 | 2.10141 | 3.15 |
| EPHX1 | 3.333333 | 0.30 |
| EREG | 3.197861 | 12.82 |
| FSCN1 | 0.407867 | 3.99 |
| ID1 | 0.30254 | 3.12 |
| KIAA1199 | 0.280992 | 0.07 |
| MBNL2 | 1.988971 | 0.33 |
| MMP1 | 9.014286 | 13.35 |
| PTGS2 | 9.625 | 6.23 |
| RARRES3 | 0.232759 | 0.22 |
| SPINK4 | 0.237762 | 0.16 |
| TNC | 3.416667 | 8.99 |

[14] Minn AJ, Gupta GP, Siegel PM, Bos PD, Shu W, Giri DD, et al. Genes that mediate breast cancer metastasis to lung. *Nature*. 2005;**436**(7050):518–524.
