## Supplementary material for "Promyelocytic Leukemia Protein regulates Angiogenesis and Epithelial-Mesenchymal Transition to limit metastasis in MDA-MB-231 breast cancer cells": primers and antibodies used in this paper

**Supplementary materials and methods**

**Primers used for qRT-PR (related to section 2.4)**

| | |
|---|---|
| PML F: 5'- GATGGCTTCGACGAGTTCAA-3' | R: 5'- GGGCTGGCTTCCTTGGATAC-3' |
| ACTIN B F: 5' CCTGTACGCCAACACAGTG 3' | R: 5'-ATACTCCTGCTTGCTGATCC 3' |
| GAPDH F:5’- CAGTCAGCCGCATCTTCTTT-3’ | R: 5'- ACCAGAGTTAAAAGCAGCCC-3’ |
| TOP2A F: 5'-TCAAACGGAATGACAAGCGA-3' | R: 5'-ATGGGCTGCAAGAGGTTTAG-3' |
| PCNA F: 5'-TTTCCTGTGCAAAAGACGGA-3' | R: 5'-CCGTTGAAGAGAGTGGAGTGG-3' |
| p21 F: 5'-CCGCTCTACATCTTCTGCCTTAGTC-3' | R: 5'-AACCTCTCATTCAACCGCCTAGTT-3' |
| HDAC9 F: 5’-GGTGGACAGTGACACCATTT-3’ | R: 5’-TGGATTCTTCAGCGTGATGG-3’ |
| MMP1 F:5’-GGGCTTTGATGTACCCTAGC-3’ | R:5’-ACTTCCGGGTAGAAGGGATT-3’ |
| CD24 F:5’-TGCTCCTACCCACGCAGATT-3’ | R:5’-GGCCAACCCAGAGTTGGAA-3’ |
| EpCAM F:5’-TTCTAAGAAAATGGACCTGACA-3’ | R:5’-TTCCCTATGCATCTCACCCA-3’ |
| CDH1 F:5’-CTCACACACCCCCTGTTGGT-3’ | R:5’-GTGAATTCGGGCTTGTTGTC-3’ |
| VEGFα F:5’-CAGAATCATCACGAAGTGGTG-3’ | R:5’-GAAGATGTCCACCAGGGTC-3 |
| CD49f (ITGA6) F: 5’-TCATGGATCTGCAAATGGAA-3’ | R: 5’- AGGGAACCAACAGCAACATC-3’ |
| IL6ST(gp130) F: 5’- CGAAGCTGTCTTAGAGTGGG-3’ | R: 5’- AAGCAAACAGGCACGACTAT-3’ |
| PIM1 F: 5’-CGAGCATACGAAGAGATCA-3’ | R: 5’-TCGGGCATCTGACAAGAGA-3’ |
| IL6 F: 5’-ACTGGCAGAAAACAACCTA-3’ | R: 5’-CAGGGGTGGTTATTGCATCT-3’ |
| TGFβR2 F: 5’-TGCCCCAGCTGTAATAGGACC-3’ | R: 5’-CCATACAGCCACACAGACTT-3’ |
| EZH2 F: 5’- TCCTTTTCATGCAACACCCA-3’ | R: 5’- TTTCAGTCCCTGCTTCCCTA-3’ |
| SLIT2 F: 5’- ACCAGTCATTTATGGCTCCTTC-3’ | R: 5’- TCAGAGAGCGTAGTCCTTGG-3’ |
| ISL1 F: 5’- GGCAATCAGATTCACGATCAG-3’ | R: 5’- GCGCATTTGATCCCGTACAA-3’ |
| XIST F: 5’- CACGTGTATGTCTCCCAGTG | R: 5'- GTGAGGCACCAATACAGAGG-3’ |
| ID1 F: 5’- AAACGTGCTGCTCTACGACA-3’ | R: 5'- GAGAATCTCCACCTTGCTCAC-3’ |
| FASCIN F: 5’- CAGCGGCCTCTCGTCTA-3’ | R: 5'- AGATCTGCTTCTTCTTCAGGC-3’ |

**Antibodies used (related to section 2.7)**

| | |
|---|---|
| PML | sc-377340 Santa Cruz, Dallas, Texas, USA |
| β-ACTIN | sc-47778, Santa Cruz |
| GAPDH | sc-32233, Santa Cruz |
| p21 | sc-397, Santa Cruz |
| STAT3 | sc-8019, Santa Cruz |
| p-STAT3 | 9145S, Cell signaling, Danvers, Massachusetts, USA |
| VIMENTIN | 5741S, Cell signaling |
| E-CADHERIN | 3195S, Cell signaling |
| GFP | sc-9996, Santa Cruz |
| ERa | M7047, DAKO, Denmark |
